# *Salmonella* hijacks a host glucose transporter for intravacuolar proliferation

**DOI:** 10.1101/2025.06.10.658782

**Authors:** Xianglan Fang, Liuliu Shi, Yan Ding, Youwei Wang, Wei Chen, Juan Xue, Long Wang, Yuanjian Hui, Xinyuan Tao, Jin Yang, Lijie Du, Duoshuang Xie, Kun Meng

## Abstract

*Salmonella* is an intracellular pathogen that resides within a vacuole, which protects it from cytosolic host defenses at the expense of limited nutrient access. Glucose serves as a critical carbon source supporting *Salmonella*’s intracellular replication. However, the molecular mechanisms driving glucose enhancement and the pathways by which cytosolic glucose becomes accessible to intravacuolar *Salmonella* remain poorly understood. Here, we elucidate a sophisticated three-pronged strategy through which *Salmonella* hijacks GLUT1 to co-opt host glucose metabolism for pathogenic advantage. Firstly, *Salmonella* infection upregulates the glucose transporter GLUT1 by activating the MAPK signaling cascade, enhancing host glucose uptake, and accelerating glycolytic flux. Secondly, *Salmonella* redirects GLUT1 to the bacterial vacuolar membrane to establish a glucose-import conduit that facilitates bacterial acquisition of cytosolic glucose. Thirdly, K29-linked ubiquitination modifications on bacterial vacuolar membranes are a previously unrecognized regulatory mechanism that potentiates GLUT1 transporter activity. Inhibition of GLUT1 potentiates *Salmonella*-triggered innate immune responses and attenuates bacterial virulence *in vitro* (RAW264.7 macrophages) and *in vivo* (murine infection model). Collectively, these findings delineate a novel paradigm of metabolic hijacking, wherein *Salmonella* systematically rewires host glucose metabolic networks to support intracellular proliferation, providing new insights into host-directed antimicrobial interventions.

## Introduction

*Salmonella enterica* is a Gram-negative facultative intracellular pathogen capable of infecting a broad range of hosts, causing diseases ranging from self-limiting gastroenteritis to systemic infections(^1, 2^). During infection, *Salmonella* survives within host cells by residing in the *Salmonella*-containing vacuole (SCV), a compartment formed through interactions with the host endolysosomal system and the subsequent formation of Salmonella-induced filaments (SIFs). These membrane structures enable immune evasion but confine the pathogen within a membrane-bound environment(^3–5^). Consequently, *Salmonella* needs to develop unique strategies to reprogram host metabolism and exploit available host metabolites to meet the bacterial energetic demands. The intracellular replication of *Salmonella* nutritionally requires the acquisition of host-derived glucose for glycolytic energy production, glycerol for membrane biosynthesis, amino acids for protein production, and GlcNAc for peptidoglycan cell wall assembly(^6, 7^). Thus, the metabolic arms race between intracellular *Salmonella* and its host dictates the outcome of infection.

Upon infection, *Salmonella* induces profound metabolic changes in host cells, most notably triggering a Warburg-like effect characterized by rapid glycolysis upregulation and oxidative phosphorylation suppression(^8^). While this metabolic shift carries potential risks for the pathogen (such as the NLRP3 inflammasome activation and enhanced phagosome acidification)(^9, 10^), *Salmonella* strategically benefits.

Increased glucose uptake and glycolytic flux may dampen some fatal immune responses. For instance, *S.* Typhimurium redirects metabolism toward glycolysis as an antioxidant defense by mitigating trans-membrane electron flow to preserve the proton motive force (ΔpH)(^11^). Concurrently, this shift augments the availability of glucose and glycolytic intermediates to fuel bacterial energy metabolism and virulence(^12^). A well-characterized paradigm involves the SPI-1 effector SopE2, which promotes the accumulation of 3-phosphoglycerate (3PG), pyruvate, and lactate.

Among these, 3PG is a direct nutrient source for intracellular proliferation, while pyruvate and lactate act as signaling molecules to activate SPI-2 virulence gene expression(^13^). Despite these insights, the molecular mechanisms underlying infection-induced glycolytic enhancement remain poorly defined. Additionally, it is still unclear what host nutrients are available to *Salmonella* within the SCV and how they enter the vacuole.

In this study, we demonstrate that *Salmonella enterica serovar* Typhimurium (*S*. Typhimurium) infection upregulates the glucose transporter GLUT1 through activation of the mitogen-activated protein kinase (MAPK) signaling cascade, leading to enhanced host cellular glucose uptake and accelerated glycolytic flux. We further show that *Salmonella* redirects GLUT1 to the bacterial-containing vacuolar membrane, establishing a putative glucose-import conduit that facilitates bacterial acquisition of cytosolic glucose. Notably, *Salmonella*-induced K29-linked ubiquitination of GLUT1 on vacuolar membranes potentiates the transporter’s activity, enhancing glucose translocation efficiency. Functional inhibition of GLUT1 not only augments *Salmonella*-triggered innate immune responses but also attenuates bacterial virulence in both *in vitro* and *in vivo* models. Collectively, these findings delineate a novel mechanism of metabolic hijacking, whereby *Salmonella* subverts host GLUT1 to reprogram glucose metabolic and immune pathways, thereby enabling its intracellular proliferation.

## Results

### *Salmonella* infection reprograms host glycolytic metabolism by upregulating GLUT1 expression

Consistent with previous findings(^12^), we demonstrated that *Salmonella*-infected macrophage cells exhibited elevated glucose uptake, as evidenced by non-radiolabeled glucose uptake and 2-NBDG fluorescence assays (Figs. 1A-B).

**Figure 1.**
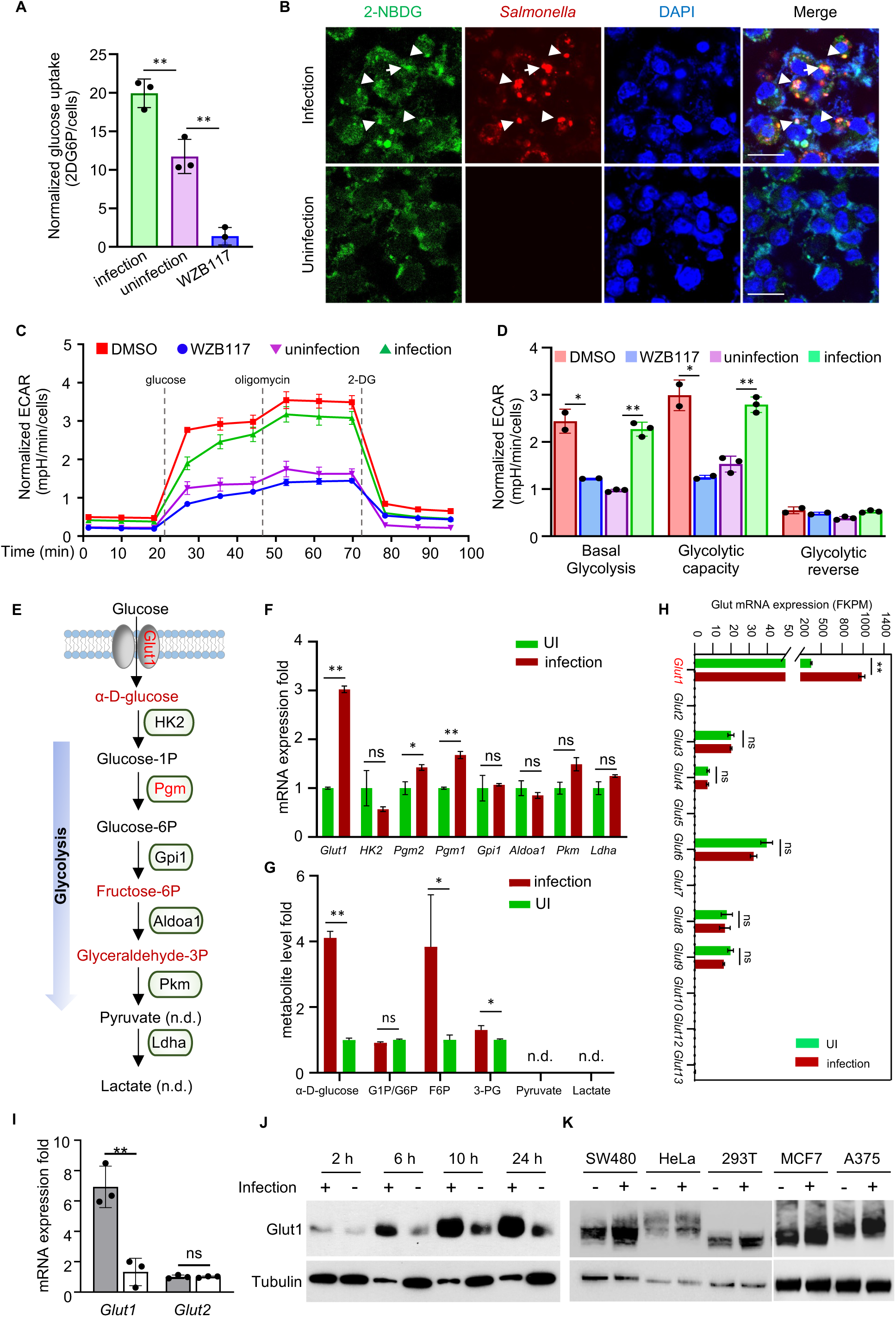
*Salmonella* infection promotes host glucose metabolism through GLUT1 upregulation. **(A-B) The effects of *Salmonella* infection on the glucose uptake of macrophages**. RAW264.7 cells were infected with *S*. Typhimurium strains for 10 h and subjected to glucose uptake assessment using either 2-DG quantification or the fluorescent glucose analog 2-NBDG imaging. (A) Quantification of intracellular 2DG6P levels normalized to total cell count. (B) Representative images showing the intensity and distribution of 2-NBDG (green) and *S*. Typhimurium (red). Scale bar, 20 μm. **(C-D) The effects of *Salmonella* infection on macrophage glycolytic activity**. RAW264.7 cells were infected with *S*. Typhimurium strains for 10 h, followed by measurement of ECAR using the Seahorse XF Glycolytic Rate Assay (C). (D) Quantitative analysis of relative ECAR levels at different glycolytic stages is presented, with the GLUT1 inhibitor WZB117 serving as the negative control. **(E-G) The effects of *Salmonella* infection on the gene and metabolite alterations in the glycolytic pathway**. (E) Schematic summary of transcriptional and metabolic alterations in the glycolytic pathway induced by *S*. Typhimurium infection. Significant changes (*p*<0.05) are highlighted in red. Non-detected metabolites are indicated as “n.d.”. (F) Transcriptomic profiling of glycolytic pathway genes in *S*. Typhimurium-infected macrophages. (G) Metabolomics profiling of glycolytic intermediates following *Salmonella* infection. **(H-K) The effects of *Salmonella* infection on GLUT1 expression in host cells**. (H) Transcriptomic analysis of Glut family member expression in *S*. Typhimurium-infected RAW264.7 macrophages. (I) qPCR validation of *Glut1* and *Glut2* mRNA expression in infected RAW264.7 cells. (J) Time-course immunoblot analysis of Glut1 protein expression in RAW264.7 cells post-infection (2-24 h.p.i.). Representative blot from three independent experiments is shown, with tubulin as the loading control. (K) Western blot analysis of GLUT1 expression across multiple cell lines at 10 h.p.i. \**p*<0.05, \*\**p*<0.01.

Intriguingly, 2-NBDG signals colocalized with intracellular *Salmonella*, implying that bacteria residing within SCV may directly utilize glucose from the host (Fig. 1B).

Seahorse extracellular flux analysis confirmed that infection enhanced glycolytic capacity, with significant increases in both basal glycolysis and glycolytic reserve (Figs. 1C-D).

To elucidate the molecular mechanisms by which *Salmonella* modulates host glucose metabolism, we tried to identify the key glycolytic components in infected RAW264.7 macrophages by transcriptomic and metabolomic analyses (Supplementary Fig. 1). *Salmonella* infection significantly upregulated the expression of *Glut1* (glucose transporter 1) and *Pgm1/2* (phosphoglucomutase 1/2), concomitant with increased intracellular levels of glucose, fructose, and 3-phosphoglycerate (3-PG) (Figs. 1E–G). Among all glucose transporters (*Glut1* to *Glut13*), Glut1 exhibited the most dramatic transcriptional induction upon *Salmonella* infection, while other Glut family members remained largely unchanged (Fig. 1H). This specific induction pattern was confirmed by qPCR analysis, which demonstrated significant upregulation of *Glut1* but not *Glut2* transcripts (Fig. 1I). Western blot analysis further demonstrated that Glut1 protein levels increased during *Salmonella* infection in a time-dependent manner (Fig. 1J). Besides, this upregulation was observed not only in RAW264.7 macrophages but also consistently across five additional cell lines (SW480, HeLa, 293T, MCF7, and A375), suggesting that *Salmonella* employs a conserved mechanism in GLUT1 regulation (Fig. 1K). Using actinomycin D (ActD) and cycloheximide (CHX) chase assays, we found that neither Glut1 mRNA stability nor protein turnover was significantly affected by *Salmonella* infection, demonstrating that Glut1 upregulation occurs primarily at the transcriptional level (Supplementary Fig. 2).

### *Salmonella* induces Glut1 expression through the Erk/Jnk-c-Jun signaling axis

To elucidate the mechanism underlying *Salmonella*-mediated GLUT1 upregulation, we performed promoter analysis of the GLUT1 gene. We identified multiple putative transcription factor binding sites, including those for MAPK (Erk1/2, Jnk, p38)-downstream effectors (c-Jun, c-Fos, CREB), as well as NF-κB p65 and HIF1α (Fig. 2A). Western blot analysis revealed that Salmonella infection specifically enhanced phosphorylation of Erk, Jnk, and their downstream target c-Jun. In contrast, activation of p38 MAPK, NF-κB p65, and HIF1α remained unchanged (Supplementary Fig. 3 and Figs. 2B-C). Pharmacological inhibition experiments demonstrated that inhibitors targeting Erk1/2, Jnk, or c-Jun significantly attenuated *Salmonella*-induced GLUT1 expression, glucose uptake activity, and GLUT1 promoter activation. In contrast, p38 and HIF1αinhibitors showed no effect (Figs. 2D-F). DNA pull-down assays further confirmed that both phosphorylated and non-phosphorylated c-Jun could directly bind to and activate the GLUT1 promoter (Figs. 2G-I), establishing a direct regulatory relationship.

**Figure 2.**
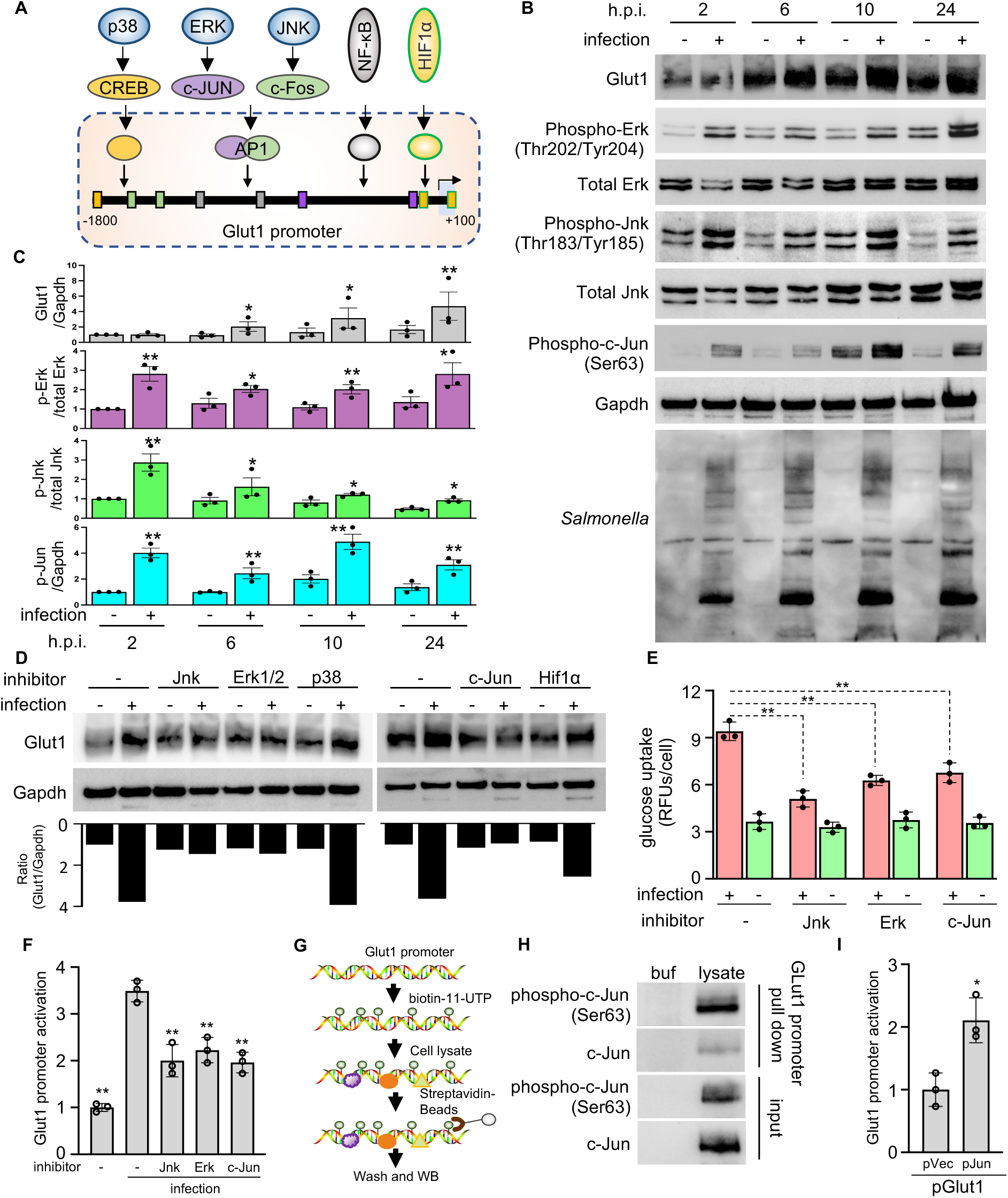
*Salmonella* infection induced Glut1 expression via Erk/Jnk-AP1 axis. **(A) *In silico* analysis of the *Glut1* promoter.** Schematic of predicted transcription factor binding sites within the Glut1 promoter region (–1800 to +100 bp relative to TSS) is shown. Potential regulatory sites are color-coded by the TF family. **(B-C) The effects of *Salmonella* infection on MAPK pathway activation. (B)** Time-course immunoblot analysis of the MAPK signaling components (Erk1/2, Jnk, p38, and c-Jun) in RAW264.7 cells post-infection (2-24 h.p.i.). Representative blots from three independent experiments are shown, with Gapdh as a loading control. Densitometric quantification of phosphorylated MAPKs was normalized to total protein (**C**). **(D-F) Pharmacological blockade of MAPK signaling attenuates *Salmonella*-induced GLUT1 upregulation and functional activity. (D-E)** RAW264.7 cells were pretreated with specific inhibitors as indicated before *Salmonella* infection and subjected to western blot or the glucose uptake assay. **(F)** 293T cells transfected with pGLUT1-Luc reporter were pretreated with MAPK inhibitors before infection. Luciferase activity was measured at 10 h.p.i. and normalized to Renilla control. **(G-I) Transcription factor c-Jun directly binds and activates the Glut1 promoter. (G)** Schematic of Glut1 promoter pull-down assay. Biotinylated DNA probes were incubated with nuclear extracts from RAW264.7 cells. **(H)** Samples were precipitated with streptavidin beads and detected with the indicated antibodies. **(I)** 293T cells were co-transfected with a pGLUT1-Luc reporter with the cJUN-plasmid and subjected to luciferase activity assay at 18 h post-transfection. \**p*<0.05, \*\**p*<0.01.

### *Salmonella* SPI-1 effectors are responsible for Glut1 expression

Lipopolysaccharide (LPS), a surface component of Gram-negative bacteria, functions as a PAMP by activating TLR4-mediated immunity extracellularly and CASP4-dependent pyroptosis intracellularly(^14, 15^). We found that various forms of LPS delivery (soluble, liposome-encapsulated, electroporated, or OMVs-delivered) failed to significantly alter Glut1 expression despite inducing marked cellular morphological changes of RAW264.7 macrophages (Supplementary Figs. 4A-B). Instead, Glut1 upregulation may represent a pathogen-specific response rather than a general host reaction to bacterial infection. Among the pathogens tested, only invasive intracellular bacteria (*Salmonella* and *Shigella*) induced significant GLUT1 expression, whereas extracellular pathogens (EPEC and *Klebsiella*) showed no such effect. Consistent with these findings, both SPI-1 T3SS-deficient (Δ*sipD*) and heat-killed (HK) *Salmonella* failed to induce Glut1 expression (Supplementary Figs. 4C-D). These results demonstrate that GLUT1 upregulation requires SPI-1 T3SS-dependent bacterial invasion. Given that *Salmonella* SPI-1 effectors (SopB/SopE/SopE2) promote invasion by hijacking host Rho GTPases for actin remodeling and simultaneously activating MAPK pathways(^16, 17^), we hypothesized that these effectors may be responsible for the upregulation of GLUT1 expression (Supplementary Fig. 4E). Indeed, ectopic expression of individual effectors in HeLa cells significantly increased Glut1 expression while synchronously activating the JNK/ERK-Jun axis (Supplementary Figs. 4F-G).

### GLUT1 interacts with membrane trafficking machinery during infection and specifically targets the vacuole-residing *Salmonella*

During early infection, nascent SCVs expand rapidly through fusion with infection-associated macropinosomes (IAMs). This process is mediated by host trafficking protein complexes, such as the SNAP25-VAMP8 pairing(^18, 19^). At later stages, SPI-2 T3SS effectors drive the biogenesis of *Salmonella*-induced filaments (SIFs) and facilitate the establishment of polarized replicative compartments (Fig. 3A).

**Figure 3.**
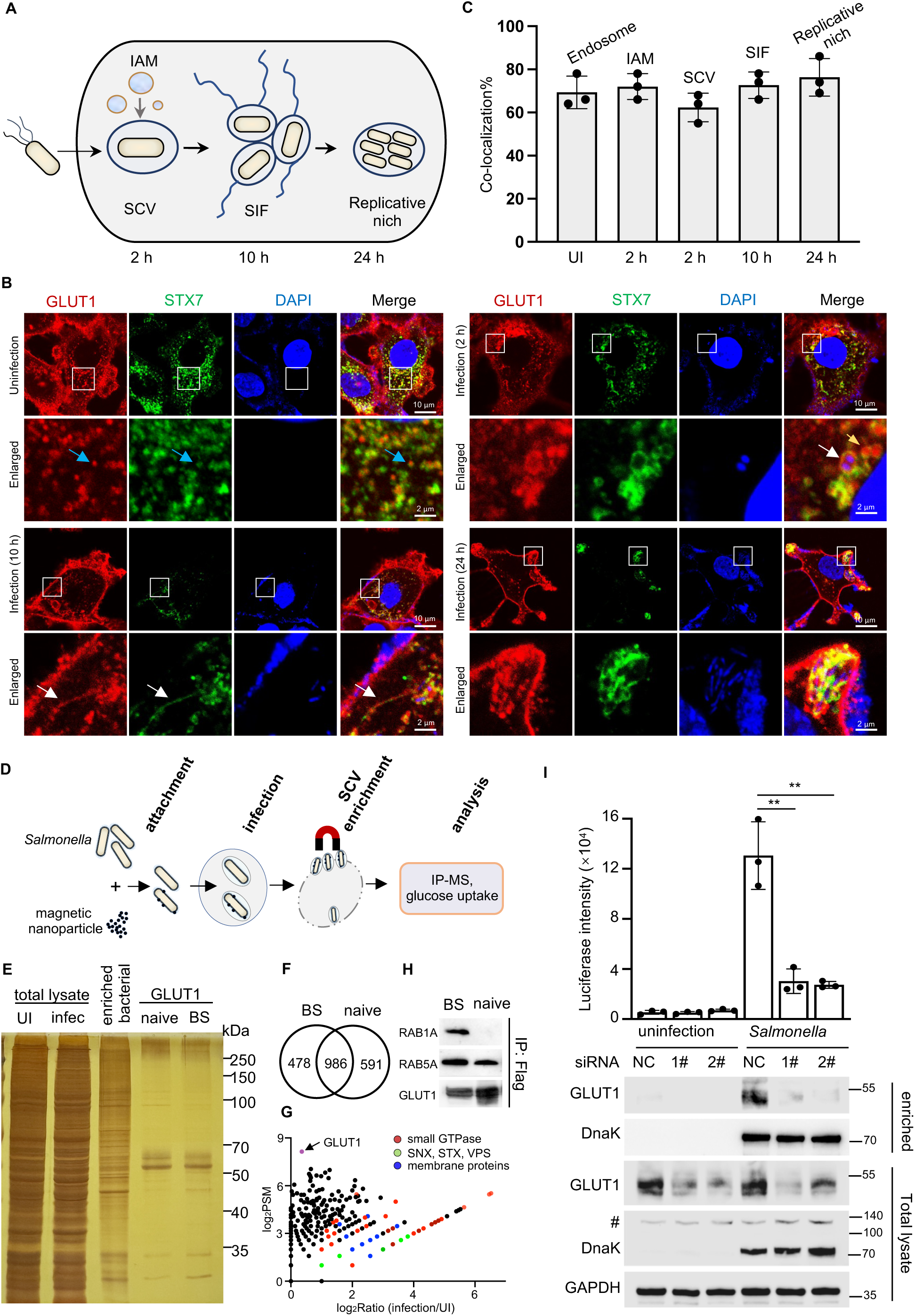
*Salmonella* redirects host Glut1 to the bacterial vacuolar membrane during intracellular infection by interacting with membrane trafficking proteins to acquire cytosolic glucose. **(A-C) The co-localization of GLUT1 with *Salmonella* intracellular vacuolar replicative structure**. (**A**) Schematic of *Salmonella’s* intracellular vacuolar structure in host cells. **(B-C)** HeLa cells stably expressing EGFP-STX7 were infected with the *S*. Typhimurium strain for the indicated time. **(B)** Representative images showing the distribution of STX7 (green), endogenous GLUT1 (red), and nucleus (blue). Scale bar, 10 µm. White boxes indicate magnified regions. Scale bar, 2 µm. Arrows indicated the co-localization of GLUT1 with endosome (blue), SCV/SIF (white), and IAM (yellow). **(C)** The co-localization rates of STX7 and GLUT1 for each sample. At least 100 cells were counted for samples from experiments done in triplicate. **(D-H) Identification of the specific GLUT1-interacting proteins on SCV. (D)** Schematic of the SCV enrichment by magnetic nanobead. **(E)** Detection of enriched proteins by silver staining. Flag-GLUT1-expressing 293T cells were either infected with *Salmonella* or left uninfected as a control. *Salmonella*-infected cells were subjected to magnetic bead enrichment of intracellular bacteria prior to anti-Flag IP. Precipitated protein complexes from both groups were separated by SDS-PAGE and visualized by silver staining. **(F-G)** Analyses of GLUT1 interactome by mass spectrometry. Proteins immunoprecipitated with an anti-Flag antibody were subjected to LC-MS/MS analysis. **(F)** Overlap analysis of GLUT1-interacting proteins identified in SCV versus uninfected cellular compartments. **(G)** Scatter plots of protein ratios as a function of their relative abundance. The ratio was calculated as spectral counts in SCV-enriched samples divided by those in uninfected samples. Large ratios indicate preferential detection in the SCV enriched sample. **(H)** Validation of GLUT1-Rab GTPase binding by immunoblotting. **(I) Effects of *Glut1* knockdown on the glucose uptake of SCV**. RAW264.7 cells were transfected with two distinct Glut1-targeting siRNA pairs for 48 h, followed by *Salmonella* infection (or mock infection) for 10 h. SCVs were isolated and subjected to glucose uptake measurement, with activity normalized to negative control (NC) siRNA-transfected cells. Glut1 knockdown efficiency was verified by immunoblotting. Data represent mean ± SD from three independent experiments (\*\**p*<0.01).

Immunofluorescence analysis revealed that, in uninfected cells, GLUT1 predominantly localized with endosomes. GLUT1 became significantly enriched on IAM and SCV membranes and maintained a persistent association with all subsequent SIFs and replicative vacuolar structures (Figs. 3B-C). Notably, GLUT1 was selectively associated with vacuole-competent pathogens (wild-type *Salmonella* and *Shigella*). Only minimal GLUT1 recruitment was observed during phagocytosis of extracellular bacteria (EPEC, *Klebsiella*) or invasion-deficient mutants (Δ*sipD* and HK *Salmonella*) by macrophage cells (Supplementary Figs. 5A-B). Additionally, when *Salmonella* ruptured the vacuolar membrane and became the cytosol super-replication status, GLUT1 colocalization was completely abolished, demonstrating its strict dependence on intact vacuolar membranes (Supplementary Figs. 5C-D).

To elucidate the potential molecular mechanism underlying GLUT1 recruitment to SCVs, we employed a nanoparticle-coated strategy for the magnetic isolation of SCVs and performed comparative interactome analysis between bacteria-associated GLUT1 (BS-GLUT1) and native GLUT1 (enriched from uninfected cells) (Figs. 3D-F). BS-GLUT1 specifically interacted with multiple host vesicular trafficking components (Fig. 3G and Supplementary Fig. 6A), including stronger binding to Rab GTPases (Rab5A/Rab1A) (Fig. 3H) and colocalization with SNARE proteins (VAMP8/Vti1b) during infection (Supplementary Figs. 6B-C). These results suggest that GLUT1 may be actively recruited to SCVs through IAM-SCV fusion by vesicular trafficking machinery proteins.

### GLUT1 is required for the glucose uptake of vacuole-residing *Salmonella*

We next investigated the functional roles of GLUT1 for the intravacuolar *Salmonella*. Despite its co-localization with SCVs and SIFs (Figure 3B), GLUT1 knockdown did not affect the integrity of these membrane structures in *Salmonella*-infected cells, as evidenced by unchanged frequencies of STX7-coated bacteria (Supplementary Figs. 7A-B). Additionally, contrary to its reported possible role in regulating autophagy(^20^), GLUT1 knockdown did not affect the bacterial xenophagy, as shown by the unchanged proportion of LC3-coated *Salmonella* Δ*sopF* mutant (Supplementary Figs. 7C-D). Instead, silencing GLUT1 led to a significant reduction in glucose uptake by the enriched SCVs (Fig. 3I). These findings suggest that GLUT1 on SCVs may serve as a critical receptor of glucose acquisition for intravacuolar *Salmonella*.

### K29-linked ubiquitination of GLUT1 on SCV enhances glucose acquisition by intravacuolar *Salmonella*

Ubiquitination modification has been reported to play roles in the stability and subcellular localization of GLUT1(^21–23^). Immunoblot analysis revealed significantly enhanced ubiquitination of BS-GLUT1, suggesting *Salmonella* infection induces additional ubiquitin modifications on GLUT1 (Fig. 4A). Quantitative mass spectrometry (MS) analysis identified four novel ubiquitinated peptides in BS-GLUT1 compared to naive GLUT1. Notably, the peptide ^232^GTADVTHDLQEMKEESR^249^ exhibited robust ubiquitination (28.7% modification rate), while other modified peptides showed less than 10% alteration. MS/MS spectral analysis precisely mapped the ubiquitination sites to K245, K255, K256, and K477 (Fig. 4B and Supplementary Figs. 8A-E). The mutation K245R/R255R/K256R/K477R (K4R) showed significantly reduced ubiquitination signals (Fig. 4C). These lysines are located within accessible GLUT1’s cytoplasmic region and evolutionary conserved among mammalian hosts of *Salmonella* (human, mouse, pig, chicken) but not across the GLUT family (Supplementary Figs. 8F-G). While this mutant retained unchanged subcellular localization in uninfected cells and SCV association in infected cells (Supplementary Fig. 9 and Fig. 4D), it significantly compromised the glucose acquisition capacity of intravacuolar *Salmonella*, resulting in reduced bacterial proliferation within SCVs (Figs. 4E-G).

**Figure 4.**
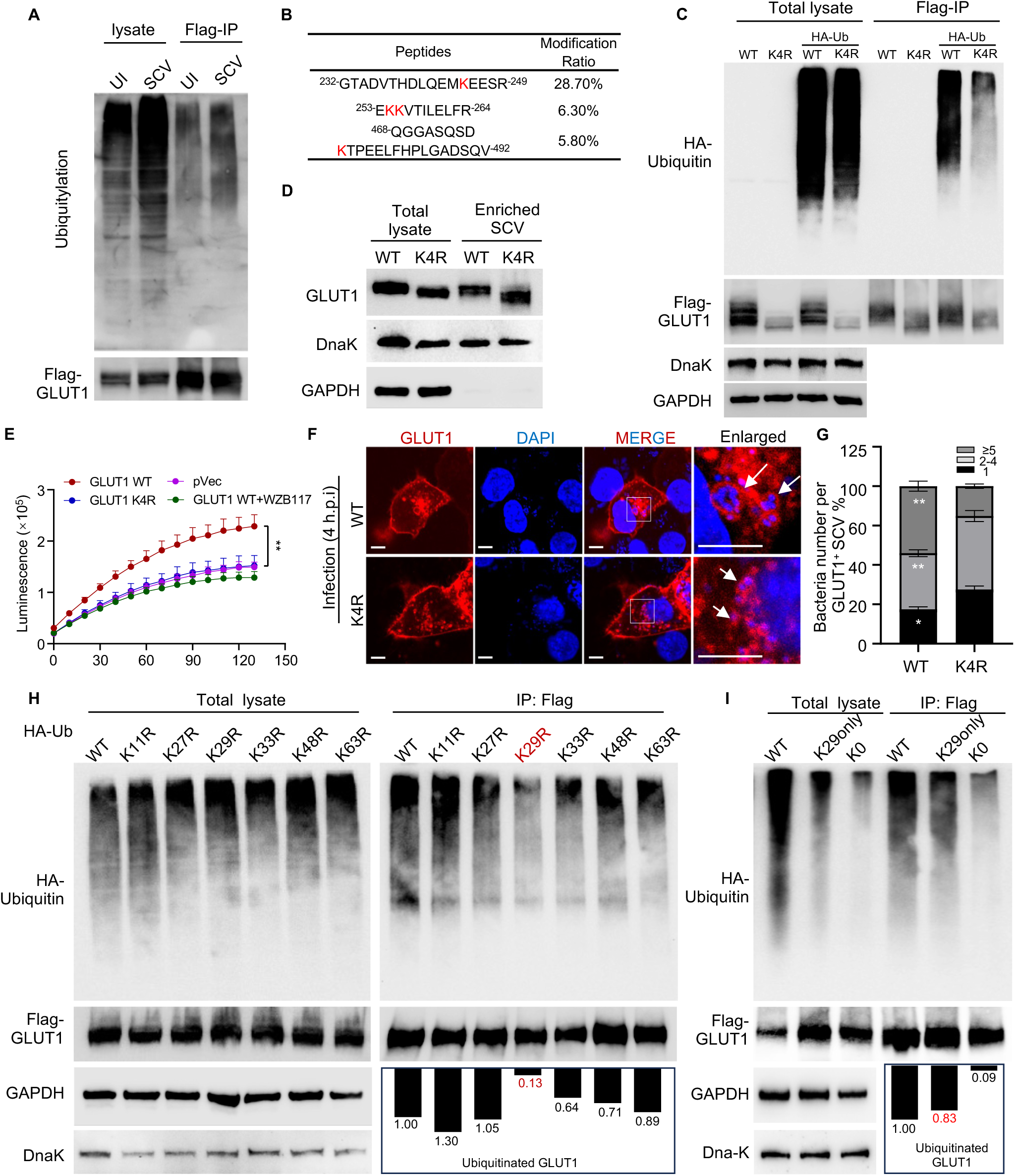
K29-linked ubiquitination of GLUT1 on SCVs facilitates glucose acquisition of the intravacuolar bacteria within host cells. **(A-C) Ubiquitination of GLUT1 on SCV occurs at K245, K255, K256, and K477. (A-B)** Parallel immunoblot and LC-MS/MS analyses were performed using the same samples as in Figure 3E. **(A)** Ubiquitin modification was detected by immunoblotting with anti-ubiquitin antibodies. **(B)** Mass spectrometry identification of GLUT1 ubiquitination sites. Modified sites are highlighted in red, with relative ubiquitination levels quantified by extracted ion chromatogram peak intensities shown in Supplementary Figure 8. **(C)** Ubiquitination analysis of GLUT1 on SCVs following lysine mutagenesis. 293T cells were co-transfected with HA-ubiquitin and either wild-type Flag-GLUT1 or its quadruple mutant K4R. At 18 h post-transfection, cells were infected with *Salmonella* for 10 h. Intracellular bacteria were magnetically enriched, followed by anti-Flag immunoprecipitation and immunoblotting. **(D-E) Effects of K4R mutation on GLUT1 recruitment to SCVs and glucose acquisition by intravacuolar *Salmonella***. HeLa cells transfected with a plasmid encoding WT Flag-GLUT1 or the K4R mutant were infected with *Salmonella* for 10 h. SCVs were magnetically isolated, followed by immunoblotting (D) and glucose uptake assay (E). For the glucose uptake assay, cells transfected with an empty vector and those treated with the GLUT1 inhibitor WZB117 were included as two distinct negative controls. **(F-G) K4R mutation reduces the bacterial numbers within SCVs. (F)** HeLa cells expressing RFP-tagged WT GLUT1 or K4R mutant were infected with *Salmonella* at MOI=10 for 4 h. Representative confocal images show the transfected GLUT1 (red) and nuclei (blue). White boxes highlight magnified regions. Arrows indicate GLUT1-positive SCVs. Scale bar: 10 μm. **(G)** Quantification of intracellular bacteria per GLUT1-positive SCV. Data from three independent experiments (more than 50 cells per group) expressed as a percentage of total GLUT1^−^positive SCVs. **(H-I) GLUT1 undergoes preferential K29-ubiquitination on SCV.** 293T cells expressing Flag-GLUT1 were co-transfected with either WT HA-ubiquitin or ubiquitin variants (single lysine mutant in **(H)**, K0, and K29 only in **(I)**) for 18 h prior to *Salmonella* infection. Intracellular bacteria were magnetically enriched, followed by anti-Flag immunoprecipitation and immunoblotting analysis with the corresponding antibodies. Box plot quantifies the relative abundance of ubiquitination (normalized to the total Flag-GLUT1). \**p*<0.05, \*\**p*<0.01.

Next, we determined the ubiquitin chain linkage type using a panel of lysine-to-arginine Ub mutants. BS-GLUT1 ubiquitination signal was most decreased in the presence of K29R or K0 (lysine-null) Ub mutants, whereas WT Ub and other single-lysine mutants (K11R, K27R, K48R, etc.) had no such significant effect. Strikingly, a K29-only Ub mutant (retaining solely K29) almost fully restored ubiquitination, indicating that BS-GLUT1 undergoes selective K29-linked polyubiquitination (Figs. 4H-I). Collectively, our data suggested that GLUT1 on SCVs facilitates glucose acquisition of the intravacuolar bacteria within host cells via its K29-conjugated ubiquitination.

### Analyses of the potential proteins catalyzing the ubiquitination of GLUT1 during

### Salmonella infection

Next, we investigated potential ubiquitin regulators for GLUT1 modification during *Salmonella* infection. Transcriptomic analysis revealed that infection significantly upregulated multiple host E3 ligases (red) and their adaptors (yellow), while downregulating several deubiquitinating enzymes (DUBs, green) (Supplementary Fig. 10A). The GLUT1 interactome demonstrated that GLUT1 complexes isolated from SCVs showed increased enrichment of four E3 ligases and reduced association with three DUBs (Supplementary Fig. 10B). Besides, *Salmonella* encoded three E3 ligases (SopA, SlrP, and SspH2)(^24^). By 10 hours post-infection, SlrP and SspH2 exhibited markedly elevated expression levels. Ectopic expression of all three ligases robustly enhanced GLUT1 ubiquitination, but SlrP-mediated modification was abrogated in the K4R mutant (Supplementary Figs. 10C-D). However, GLUT1 ubiquitination on SCVs may represent a multifaceted regulatory process that necessitates in-depth mechanistic exploration.

### GLUT1 inhibition restricts *Salmonella* intracellular replication by enhancing host innate immune responses and disrupting bacterial metabolic pathways

To investigate the role of GLUT1 in *Salmonella* replication, we first silenced GLUT1/Glut1 expression using two effective siRNA pairs in SW480 intestinal epithelial cells and RAW264.7 macrophages (Fig. 5A). While GLUT1/Glut1 knockdown did not affect bacterial invasion (1 h post-infection), it significantly enhanced *Salmonella* replication at 24 h in both cell types (Fig. 5B). This phenotype was recapitulated using the GLUT1-specific inhibitor WZB117, which suppressed bacterial proliferation in macrophages without affecting growth in LB medium (Fig. 5C-E), establishing GLUT1 as a key restriction factor for vacuolar *Salmonella* replication.

**Figure 5.**
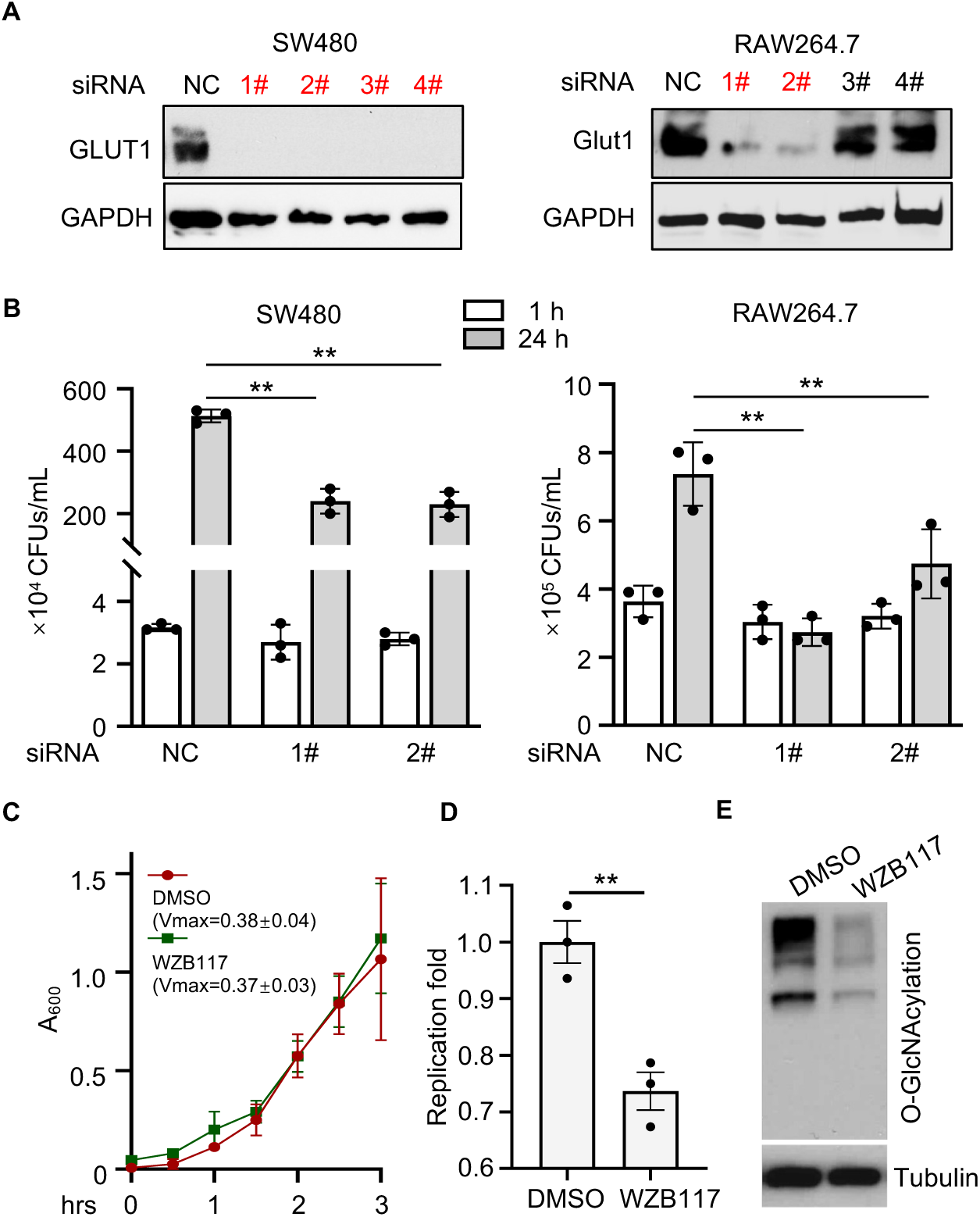
GLUT1 is essential for efficient *Salmonella* infection in host cells. **(A-B) The effects of GLUT1 knockdown on the intracellular replication of *Salmonella.*** RAW264.7 and SW480 cells were transfected with *Glut1*/*GLUT1*-targeting siRNA pairs for 48 h, followed by *Salmonella* infection at a multiplicity of infection of 10. Fold replication was determined by comparing bacterial counts at 2 and 24 h post infection. **(A)** Validation of GLUT1 silencing efficiency by siRNA. **(B)** Quantification of intracellular bacterial loads by serial dilution plating assay. **(D) Effect of WZB117 treatment on *Salmonella* growth kinetics in LB broth**. **(E-F) Effects of WZB117 treatment on the replication of *Salmonella* in macrophages**. (**E**) RAW264.7 cells were infected with *Salmonella* for 2 h, followed by treatment with the GLUT1 inhibitor WZB117. At 24 h post-infection, cells were lysed, and intracellular bacterial loads were quantified. **(F)** Validation of GLUT1 inhibitor efficacy through *O*-GlcNAcylation status analysis by immunoblotting. Results shown are mean values ± SD (error bar) from three independent experiments. \*\**p*<0.01.

To elucidate the functional consequences of GLUT1 inhibition during *Salmonella* infection, we performed dual RNA-seq to simultaneously analyze host and bacterial transcriptomes (Fig. 6A). Notably, KEGG and GO analyses of differentially expressed genes revealed that GLUT1 suppression via WZB117 broadly inhibited host metabolic pathways (green) while activating innate immune responses (red), including enhanced expression of pro-inflammatory cytokines (Figs. 6B-D). These transcriptomic findings were subsequently validated by qPCR and ELISA assays (Figs. 6E-F). Parallel bacterial profiling showed significant alterations in *Salmonella* metabolic pathways, particularly central carbon metabolism, amino acid metabolism, membrane transport, and two-component regulatory systems (Figs. 6G-I). These findings demonstrate that GLUT1 serves as both a critical metabolic hub supporting *Salmonella* intracellular replication and a key suppressor of host anti-*Salmonella* immunity.

**Figure 6.**
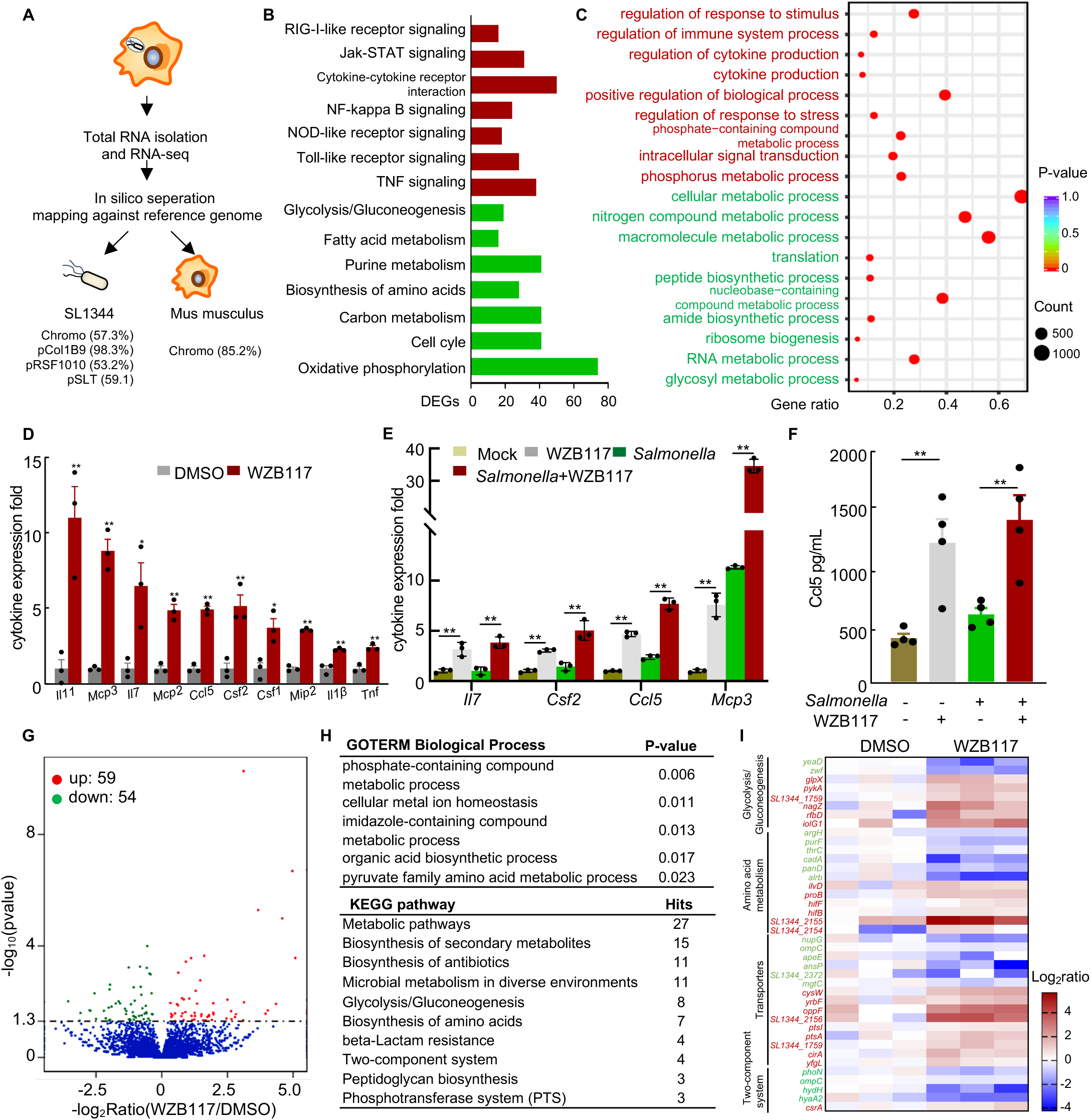
Transcriptional reprogramming during *Salmonella* infection under GLUT1 inhibition. RAW264.7 macrophages were infected with *S*. Typhimurium for 2 h, followed by treatment with 50 μM WZB117 for an additional 10 h prior to analyses. **(A) Schematic of dual RNA-seq strategy in *Salmonella*-infected macrophages**. Read mapping statistics shows the genomic and plasmid coverage distribution. **(B-C) Host Transcriptomic Responses to GLUT1 Inhibition during Infection.** Pathway enrichment analyses of DEGs were performed via Gene Ontology **(B)** and KEGG **(C)** using the DAVID online tool. The bar length and circle size denote the number of DEGs enriched in each pathway. Red labels indicate upregulated pathways, while green labels indicate downregulated pathways. **(D-F) Effects of GLUT1 inhibition on cytokine production during *Salmonella* infection. (D)** Transcriptomic profiling of cytokine expression during *Salmonella* infection. **(E)** qPCR validation of cytokine mRNA levels. **(F)** ELISA quantification of secreted cytokine proteins. \**p*<0.05, \*\**p*<0.01. **(G-I) Effects of GLUT1 inhibition on *Salmonella* gene expression. (G)** Volcano plot analysis of differentially expressed bacterial genes. **(H)** Functional enrichment analysis of transcriptional changes of *Salmonella*. **(I)** Hierarchical clustering heatmap of bacterial differentially expressed genes involved in the major biological process. Red labels correspond to the upregulated DEGs, and green labels correspond to the downregulated DEGs.

### Glut1 inhibition enhances *S.* Typhimurium-induced immune responses and reduces bacterial virulence *in vivo*

Next, we investigated the role of Glut1 in *Salmonella* infection using a murine model (Figs. 7A-B). Infection significantly upregulated Glut1 expression in the liver and spleen, as validated by qPCR, immunoblotting, and immunohistochemistry (Figs. 7C-F). Pretreatment with the GLUT1 inhibitor WZB117 significantly improved survival outcomes (Fig. 7B), and attenuated *Salmonella*-induced pathological damage, with decreased necrotic lesions of the liver and splenomegaly (Figs. 7G-I). This protective effect correlated with a marked reduction in bacterial burden, as evidenced by decreased *Salmonella* colonization in the liver and spleen (Fig. 7J). Glut1 inhibition enhanced host defense responses, as indicated by elevated *Il6* mRNA levels in infected tissues (Fig. 7K) and increased serum concentrations of chemokines (Fig. 7L). These findings demonstrate that GLUT1 inhibition protects against *Salmonella* infection by potentiating host immune responses.

**Figure 7.**
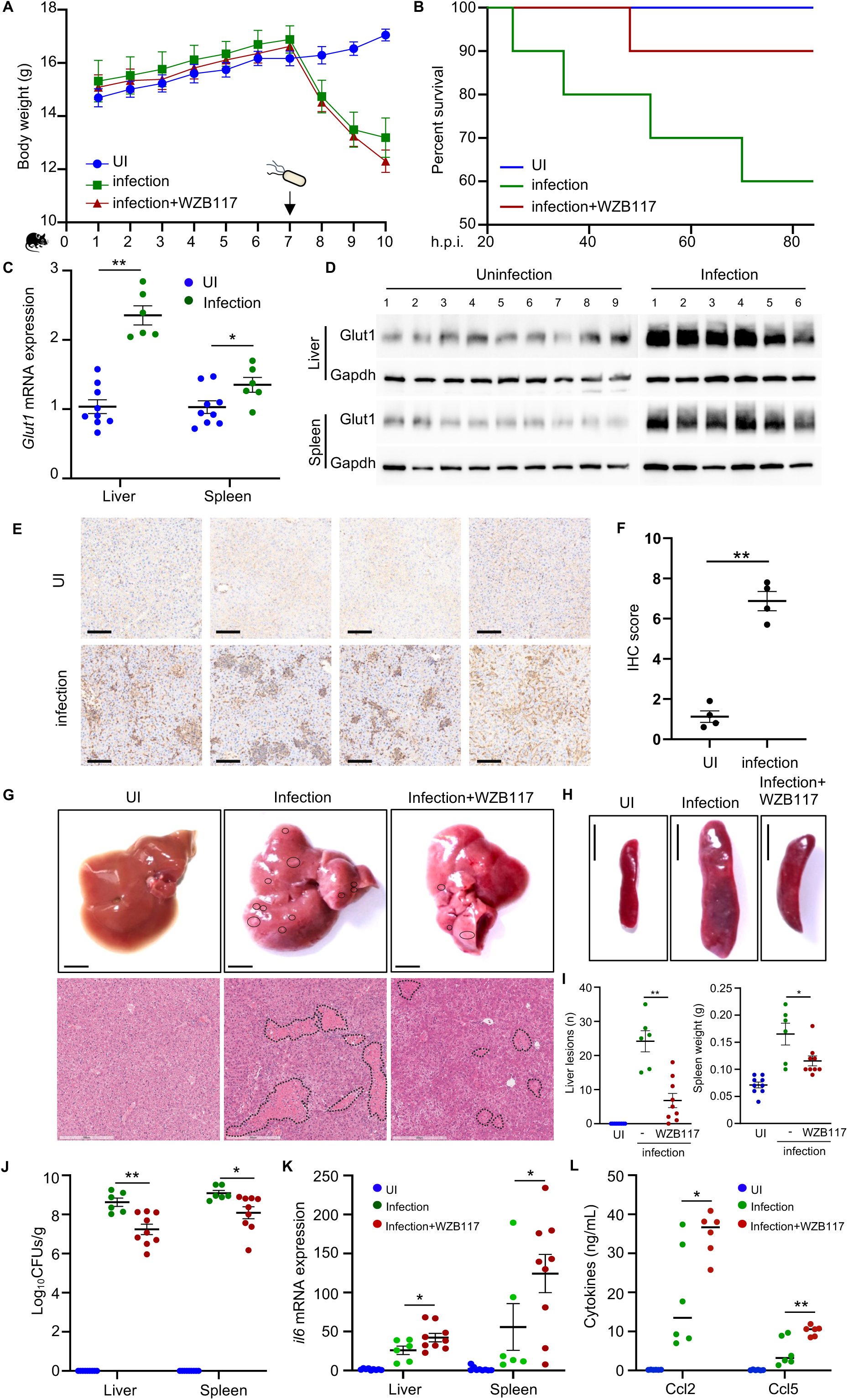
Glut1 inhibition enhances *S.*Typhimurium-induced immune responses and reduces bacterial colonization *in vivo.* **(A) Experimental procedure of *Salmonella* infection and pharmacological intervention**. C57BL/6 mice (n=10 per group) were received daily intraperitoneal injections of WZB117 for 7 consecutive days before *Salmonella* infection and subjected to the following analyses. Body weight was monitored daily throughout the experimental period. **(B) Kaplan-Meier survival curves of *Salmonella*-infected mice for 82 hours post-infection**. **(C-F) The effects of *Salmonella* infection on the Glut1 expression in the liver and spleen**. *Glut1* mRNA levels were examined by qPCR (C), and the Glut1 protein expression levels were examined by immunoblots (D) or immunohistochemistry (E-F). **(G-I) The effects of Glut1-inhibition on the *Salmonella*-induced hepatic and splenic pathology. (G)** Shown are the representative images of liver gross morphology (top; scale bar, 2 mm) and responding H&E staining (bottom; scale bar, 300 µm). Boxed areas indicate necrotic lesions. **(H)** Shown are representative images of spleen gross morphology. **(I)** Quantification of the hepatic and splenic pathological damage. **(J) The effects of Glut1 inhibition on the bacterial colonization.** Bacterial counts in the liver and spleen were calculated as colony-forming units (CFUs) per gram of tissue. **(K-L) The effects of Glut1-inhibition on the inflammatory response in *Salmonella*-infected mice**. (I) *Il6* mRNA levels were quantified by qPCR. Serum concentrations of Ccl2 and Ccl5 were measured by ELISA. \**p*<0.05, \*\**p*<0.01.

## Discussion

The strategy whereby intracellular pathogens co-opt host nutrient receptors for survival epitomizes the evolutionary arms race between microbes and their hosts. Our study demonstrates an unrecognized mechanism by which *Salmonella* ensures intracellular survival through coordinated regulation of host GLUT1. First, the SPI-1 T3SS upregulates GLUT1 expression via MAPK signaling, enhancing glucose uptake to fuel bacterial replication. Second, SPI-1 T3SS also mediates GLUT1 recruitment to the SCV membrane. Concurrently, *Salmonella* induces K29-linked ubiquitination of GLUT1, further enhancing its glucose transport activity. GLUT1 inhibition not only disrupts *Salmonella*’s metabolic adaptation but also potently enhances anti-*Salmonella* innate immune responses, highlighting GLUT1 as both a metabolic vulnerability and an immunoregulatory node during infection. These findings establish a new paradigm in microbial pathogenesis, illustrating how an intracellular pathogen can simultaneously manipulate host biology across multiple levels—transcriptional regulation, spatial protein trafficking, and activity modulation through post-translational modifications (PTMs)—to optimize its intracellular niche (Fig. 8).

**Figure 8.**
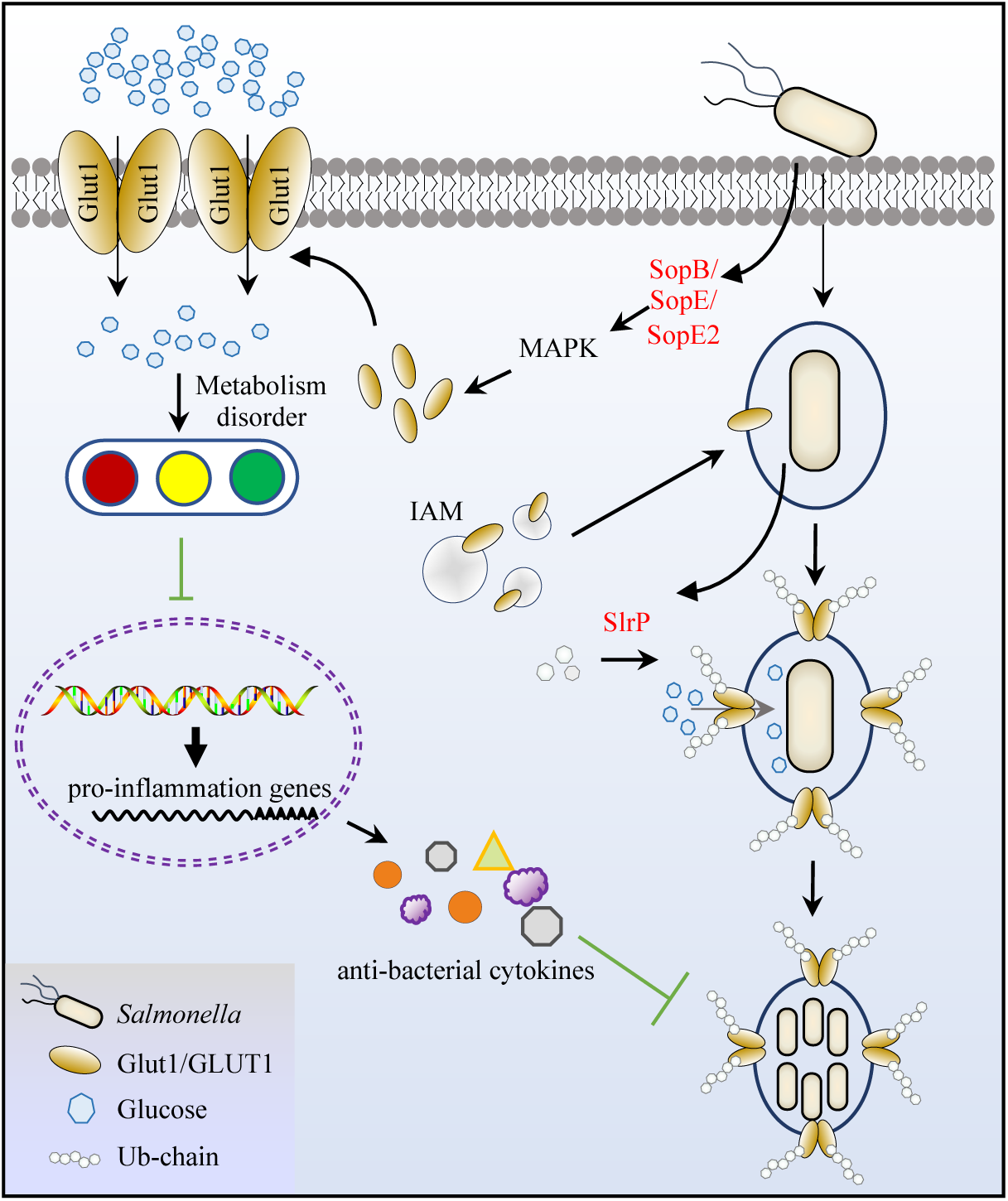
The schematic diagram of this work.

### Novel SPI-1-MAPK regulatory mechanism of GLUT1 expression during

### Salmonella infection

The expression of GLUT1 exhibits exquisite sensitivity to environmental stressors, including nutrient deprivation, hypoxia, and LPS stimulation(^25–27^). This regulation is governed by a sophisticated network of molecular mechanisms: transcriptional activation via binding of factors (e.g., NF-κB and HIF-1α) to the GLUT1 promoter(^28, 29^), post-transcriptional modulation through miRNA-mediated mRNA destabilization (e.g., miR-320a/miR-328-3p interaction with the UTR region via RNA-induced silencing complex (RISC)(^30, 31^), and post-translational control of protein turnover via ubiquitination-mediated proteolysis facilitated by specific E3 ligases(^22, 23^).

In this study, we show that GLUT1 expression is upregulated through a mechanism distinct from canonical pathways. During *Salmonella* infection, neither mRNA stability nor protein degradation of GLUT1 is altered, and this upregulation is independent of LPS stimulation or direct regulation by HIF-1α/NF-κB. Instead, this process depends on the SPI-1 and involves a previously unrecognized MAPK-dependent regulatory mechanism. Overexpression of the SPI-1 T3SS effectors SopB/E/E2, beyond their known redundant roles in Rho GTPase-mediated invasion, significantly enhances GLUT1 levels while concurrently activating MAPK signaling cascades. Thus, our findings expand the paradigm of these effectors beyond cytoskeletal manipulation to direct metabolic regulation.

### GLUT1 recruitment and ubiquitination induced by *Salmonella* for vacuolar nutrient supply

Although pathogens residing within membrane-bound vacuoles can evade cytosolic antimicrobial peptides and immune responses, this isolation also cuts off their access to host nutrients. Consequently, pathogens must actively hijack host nutrient transport systems to sustain their survival and replication. While extensive research has focused on pathogen interactions with the host endolysosomal system (e.g., RAB proteins, SNAREs) to establish and stabilize replication niches(^19, 32^), the mechanisms by which intracellular pathogens selectively recruit host nutrient transporters remain poorly understood.

To our knowledge, only a few studies have documented such phenomena. *Legionella* recruits host SLC1A5 to the *Legionella*-containing vacuole (LCV) to facilitate amino acid uptake(^33^). *Chlamydia* co-opts the host transporter SLC35D2 to import UDP-glucose into the vacuole for bacterial glycogen synthesis(^34^). Although previous studies have shown that *Salmonella* can recruit host transporter proteins SLC1A5, SLC3A2, and SLC7A5 to the SCV membrane to subsequently mobilize mTOR and evade host autophagy(^35^), whether these transporters directly contribute to nutrient acquisition by intracellular pathogens remains unclear. In this study, we uncovered a novel mechanism by which *Salmonella* specifically recruits the glucose transporter GLUT1 to the SCV membrane. Our findings demonstrate that intracellular bacteria can access the host glucose by using the fluorescent glucose analog 2-NBDG, and more importantly, GLUT1 knockdown severely impairs SCV glucose import, establishing the critical role of GLUT1 recruitment by *Salmonella* for its intravacuolar nutrient acquisition. Notably, GLUT1 localizes to all intracellular replicative compartments, indicating an important role in systemic nutrient scavenging during intravacuolar lifestyle.

The mechanism of GLUT1 recruitment appears SPI-1 T3SS-dependent, as phagocytosed bacteria (HK/Δ*sipD* Salmonella, EPEC, Klebsiella) show impaired GLUT1 decoration. Consistently, GLUT1 associates with IAM, whose formation relies on SPI-1 T3SS-mediated cytoskeletal rearrangement during invasion, linking T3SS-dependent invasion to GLUT1 recruitment. Given the role of IAM-SCV fusion in facilitating SCV expansion and the observed colocalization of GLUT1 with multiple vesicular transport proteins involved in fusion, we propose that GLUT1 is incorporated into the SCV through IAM-SCV membrane fusion events following *Salmonella* invasion. However, the exact relationship requires further investigation, particularly since inhibition of IAM-SCV fusion frequently disrupts normal SCV biogenesis.

While GLUT1 ubiquitination has been previously shown to modulate its stability or intracellular trafficking, we identify a distinct pathogen-driven mechanism in which *Salmonella* specifically induces K29-linked ubiquitination of SCV-recruited GLUT1. Notably, this modification targets lysine residues in cytosolic loops, enhancing glucose transport activity without altering protein turnover or localization. The precise mechanism underlying the *Salmonella*-induced GLUT1 ubiquitination remains challenging to define due to the global reprogramming of the host ubiquitin system during infection and the transient (“kiss-and-run”) nature of ubiquitin enzyme-substrate interactions. We identified *Salmonella*-encoded E3 ligase SlrP as capable of specifically modifying GLUT1 at these ubiquitination sites, suggesting its potential involvement. However, more refined systems are required for definitive validation.

Collectively, our findings demonstrate that *Salmonella* has evolved a dual strategy (recruitment and activity enhancing PTM of GLUT1) for intravacuolar nutritional support.

### *Salmonella* hijacks GLUT1 to evade host immune defenses

GLUT1 is well-established as a proinflammatory regulator that modulates both innate and adaptive immune responses. In innate immunity, it promotes macrophage polarization toward the M1 phenotype, activates NF-κB signaling, and stimulates the production of proinflammatory cytokines including TNF-α, IL-6, and IL-1β(^36^).

Within adaptive immunity, GLUT1 enhances T cell activation and effector functions (such as IFN-γ and TNF-α secretion) while facilitating B cell differentiation into antibody-producing plasma cells(^37, 38^).

The precise mechanisms underlying GLUT1’s regulation of anti-pathogen immunity, especially against intracellular pathogens, remain elusive due to the intricate nature of host-pathogen interactions. GLUT1 exacerbates lung fibrosis through AIM2 inflammasome activation during *Streptococcus pneumoniae* infection and is essential for monocyte inflammatory responses via HIF-1α-mediated glycolysis in *Treponema pallidum* infection(^39^). In contrast to these findings, our study reveals a novel anti-inflammatory role of GLUT1 during *Salmonella* infection, where its inhibition in both macrophages and mice unexpectedly enhances inflammatory responses with increased cytokine expression and secretion. We propose that GLUT1-driven glycolysis generates key immunometabolites that suppress innate immunity during infection. Specifically, elevated glucose levels may promote UDP-GlcNAc production via the hexosamine biosynthetic pathway (HBP), potentially contributing to immune suppression. Host O-GlcNAc transferase (OGT) utilizes UDP-GlcNAc to modify critical immune regulators like NF-κB subunits and RIPK3 through O-GlcNAcylation, thereby dampening inflammatory signaling(^40^). Concurrently, Salmonella effector proteins (SseKs) exploit host UDP-GlcNAc to glycosylate death receptors (TNFR1 and Fas) via Arg-GlcNAcylation, further blocking immune activation(^41, 42^). Additionally, infection-induced accumulation of pyruvate and lactate stimulates expression of the SPI-2 virulence system required for the secretion of effector proteins that comprehensively impair host innate immunity(^13^). In this regard, *Salmonella* infection upregulates GLUT1 to create an immunosuppressive microenvironment that favors bacterial survival and proliferation.

## Materials and methods

### Plasmids, antibodies, and reagents

For transient expression studies, the full-length open reading frames (ORFs) of GLUT1 and c-JUN were amplified from a HeLa cDNA library. *Salmonella* effector genes (SopB, SopE, SopE2, SopA, SlrP, and SspH2) were amplified from the genomic DNA of *Salmonella* Typhimurium strain SL1344. These fragments were subsequently cloned into the pCS2-EGFP, pCS2-RFP, or pCS2-Flag vectors for mammalian cell expression. The plasmids pRK5-HA-Ub-WT and its lysine mutants were maintained in our laboratory(^43^). The reporter plasmids pGLUT1-Luc and pRL-TK were commercially obtained from MiaoLing Biology Company. The GLUT1 K4R mutant was generated by site-directed mutagenesis using a standard PCR-based cloning strategy. All constructs were verified by DNA sequencing prior to experimental use.

The following commercial antibodies were listed in this study. Antibodies for Glut1 (E4S7I, #73015), NF-κB p65 (D14E12, #8242), phospho-NF-κB p65 (Ser536, 93H1, #3033), Erk1/2 (137F5, #4695), phospho-Erk1/2 (Thr202/Tyr204, D13.14.4E, #4370), JNK (#9252), phospho-JNK (Thr183/Tyr185, 81E11, #4668), p38 (D13E1, #8690), phospho-p38 (Thr180/Tyr182, D3F9, #4511), c-Jun (60A8, #9165), phospho-c-Jun (Ser73, D47G9, #3270), and HIF1α (D1S7W, #36169S) were from Cell Signaling Technology. Antibodies for Flag (AE092), HA (AE105), GFP (AE078), GAPDH (A19056), β-Actin (AC026), α-Tubulin (AC012), and H3 (A22348) were purchased from Abclonal. Antibodies for *Salmonella* (ab35156), DnaK (ab69617), and O-GlcNAc (ab2739) were from Abcam, and anti-Rab1 (11671-1-AP) and anti-Rab5 (11947-1-AP) antibodies were from Proteintech. The GLUT1 inhibitor WZB117 (T7018) was purchased from TargetMol. All cell culture reagents were obtained from Invitrogen, and other chemicals were from Sigma-Aldrich unless otherwise noted.

### Cell culture and transfection

HeLa (RRID: CVCL_0030), 293T (RRID: CVCL_0063), RAW264.7 (RRID: CVCL_0493), MCF7 (RRID: CVCL_0031), SW480 (RRID: CVCL_0546), and A375 (RRID: CVCL_0132) cell lines were obtained from the American Type Culture Collection (ATCC). Stable HeLa cell lines expressing EGFP or EGFP-tagged proteins (VAMP8, STX7, VTI1B, SNAP25, or LC3) were generated in our previous studies and maintained in our laboratory(^19^). All cells were cultured in high-glucose Dulbecco’s modified Eagle medium (DMEM; HyClone) supplemented with 10% fetal bovine serum (FBS; Gibco), 2 mM L-glutamine, and 1% penicillin/streptomycin at 37°C in a humidified 5% CO2 atmosphere. All cell lines exhibited consistent growth characteristics and demonstrated the expected biological responses, including stable bacterial infection efficiency, high transfection efficiency, and inflammatory responses.

For transient transfection, cells were transfected using JetPrime transfection reagent (Polyplus) following the manufacturer’s protocol. For siRNA-mediated gene silencing, 2 × 10⁶ cells were transfected with 200 pmol of siRNA using the following sequences: mGlut1 #1 (5’-GGACCUCAAACUUCAUUGUTT-3’), mGlut1 #2 (5’-CUGUCGGCCUCUUUGUUAATT-3’), hGLUT1 #1 (5’-CAGCCAAAGUGACAAG ACATT-3’), hGLUT1 #2 (5’-CCAAGAGUGUGCUAAAGAATT-3’), or negative control siRNA (5’-TTCTCCGAACGTGTCACGT-3’). Luciferase activity was assessed using the Dual-Luciferase Reporter Assay System (Promega) according to the manufacturer’s instructions.

### Bacterial culture and infection

The pathogenic bacterial strains (*Salmonella* Typhimurium SL1344, EPEC E2348/69, *Shigella flexneri* 2457T, *Klebsiella pneumoniae* JH1) were maintained in our lab. *Salmonella* Typhi Ty2 was kindly provided by Prof. Lu Feng and Prof. Lingyan Jiang of Nankai University. The SPI T3SS mutant derivatives of *S*. Typhimurium Δ*sipD* and Δ*ssaV* were generated by a standard homologous recombination method using the suicide plasmid pCVD442 in our previous study(^32^). Bacteria were cultured in LB Broth at 37 °C with shaking. When necessary, cultures were supplemented with antibiotics with the following final concentrations: streptomycin, 100 μg mL^−1^; ampicillin, 100 μg mL^−1^; kanamycin, 50 μg mL^−1^.

Bacterial infection of mammalian cells followed a previously described protocol with a few modifications(^32^). Briefly, pathogenic bacteria were cultured overnight (approximately 16 h) at 37 °C on a shaker (220 rpm/min) and then subcultured in antibiotic-free LB at a dilution of 1:33 for another 3 h. Infection was performed at a multiplicity of infection (MOI) of 100 for 30 min at 37 °C. To promote and synchronize infection, 24-well plates were centrifuged at 700 *g* for 5 min at room temperature. A gentamycin-killing assay was performed to remove and kill the extracellular bacteria. Briefly, cells were washed with PBS twice, and the culture medium was replaced with fresh medium supplemented with 100 μg mL^−1^ gentamycin. After a further 1.5 h incubation at 37 °C, 5% CO2, the supernatant was replaced with a medium containing 20 μg mL^−1^ gentamicin. After incubation for an indicated period, infected cells were collected and subjected to further immunoprecipitation or immunofluorescence.

To measure *S*. Typhimurium replication within host cells, SW480 and RAW264.7 cells were infected with the indicated *Salmonella* strains with an MOI of 10. A gentamycin-killing assay was performed as described above. At 2 h and 24 h post-infection, cells were lysed in cold PBS containing 1% Triton X-100, and serial dilutions were plated on the appropriate antibiotic agar plates to determine the colony-forming units. The replication fold was the number of intracellular bacteria at 24 h divided by the number at 2 h.

### Mice infection and S. Typhimurium virulence assays

Five to six-week-old wild-type C57BL/6 mice were purchased from the Experimental Animal Center of Hubei University of Medicine (Hubei, China) and were maintained in the specific pathogen-free (SPF) facility at Hubei University of Medicine. All procedures were approved by the Institutional Animal Care and Use Committee at Hubei University of Medicine (#03125020K) and conducted in compliance with national animal welfare guidelines.

C57BL/6 mice (n=10 per group) were pretreated with daily intraperitoneal injections of WZB117 (10 mg kg^−1^) or vehicle control for 7 days prior to intraperitoneal injections with 5×10⁴ CFU of *S*. Typhimurium. Mortality was monitored for 82 h, after which surviving mice were euthanized for tissue collection. Serum was collected for cytokine ELISA. Liver and spleen samples were processed for bacterial enumeration (serial dilution plating), histopathology (H&E staining), and molecular analyses (WB, RT-qPCR). Statistical significance (*p*<0.05) was determined using unpaired *t-*tests in GraphPad Prism.

### Actinomycin D and cycloheximide chase assays

For mRNA stability assessment, cells were treated with 1 μg mL^−1^ actinomycin D to block transcription, followed by RNA extraction at 0-10 h for RT-qPCR analysis.

mRNA levels were normalized to *Gapdh* and expressed as relative quantities compared to 0 h. For protein degradation analysis, cells were treated with 100 μg mL^−1^ cycloheximide to inhibit new protein synthesis, and then harvested at the indicated time for western blotting. Protein levels were normalized to Tubulin and calculated as percentages relative to 0 h. All experiments were performed with triplicate biological replicates, and the degradation rates were determined by plotting relative levels at each time point.

### Magnetic nanoparticle labeling of *Salmonella* and SCV isolation

*Salmonella* labeling and SCV isolation were performed as previously described with modifications(^44^). Briefly, carboxyl-coated paramagnetic Fe₃O₄ nanoparticles (30 nm diameter; Zhongkekeyou Beads, China) were prepared by sonicating an aliquot of the 3 mg/mL stock solution to disperse aggregates, followed by 1:10 dilution in sterile deionized water. Mid-log phase *Salmonella* cultures (OD₆₀₀ ≈ 0.8) were pelleted (16,000 g, 5 min), resuspended in PBS, and incubated with nanoparticles at a 5:1 (v/v) ratio (37°C, 20 min, 200 rpm shaking). Labeled bacteria were purified through a 0.2 μm syringe filter, washed twice with PBS, and recovered by back-flushing with 1 mL PBS. Bacterial viability and concentration were determined by serial dilution plating on LB agar. Bacterial concentration was determined by serial dilution plating.

For SCV isolation, mammalian cells were infected with labeled *Salmonella* at MOI= 50 for 10 h. Cells were washed with PBS and lysed in 0.1% Triton X-100. Lysates were subjected to magnetic separation (Magnetic Separation Rack, NEB) for 5 min at 4°C. The magnetic fraction was washed twice with PBS containing 2% sucrose and finally resuspended in 100 μL of the same buffer. Isolated SCVs were either processed immediately or stored at –80°C for downstream applications including glucose uptake assays, immunoprecipitation, or immunoblotting.

### Glucose uptake and seahorse analysis

Glucose uptake was quantified using the Glucose Uptake-Glo™ Assay (Promega) following the manufacturer’s protocol with some modifications. Briefly, SCV-enriched particles or intact cells were washed with PBS and incubated with 1 mM 2-deoxyglucose (2-DG) in PBS for 10 min. The reaction was terminated using acidic Stop Buffer, followed by neutralization. Lysates were then mixed with 2DG6P detection reagent and incubated for 1 h at room temperature. Luminescence was quantified using a microplate reader to determine 2DG6P levels. Alternatively, RAW264.7 macrophages grown on coverslips were mock-infected or infected with *Salmonella* for 8 h, followed by incubation with 100 μg mL^−1^ fluorescent 2-NBDG in glucose-free DMEM for 30 min at 37°C. After fixation, intracellular 2-NBDG localization and fluorescence intensity were analyzed by confocal microscopy.

The extracellular acidification rate (ECAR) was measured using the Seahorse XF Glycolytic Rate Assay Kit (Agilent Technologies) following the manufacturer’s protocol. Briefly, RAW264.7 macrophages were mock-infected or infected with Salmonella for 10 h, then seeded onto Seahorse XF96 cell culture microplates at a density of 2 × 10⁴ cells per well. Prior to the assay, cells were washed and maintained in Seahorse XF DMEM medium (pH 7.4) supplemented with 2 mM glutamine, 10 mM glucose, and 1 mM pyruvate. The assay was performed using a Seahorse XFe96 Analyzer with the following measurement cycle: 3 min mixing, 2 min waiting, and 3 min recording. After baseline ECAR measurements, sequential injections of 10 μM oligomycin and 50 mM 2-DG were performed to assess glycolytic flux. Data were normalized to total cell number and analyzed using the Wave Desktop Software (Agilent Technologies).

### Biotinylated DNA synthesis and DNA pull-down assay

The Glut1 promoter region (–1800 to +100 bp relative to TSS) was PCR-amplified from RAW264.7 genomic DNA using a biotin-11-UTP: TTP mixture (1:10 ratio) to generate the biotinylated DNA. Successful biotin labeling was confirmed by electrophoretic detection, with labeled fragments exhibiting slight size retardation compared to unmodified DNA. For DNA pull-down assay, 2 μg of biotinylated probes were incubated with 500 μg of RAW264.7 nuclear extracts in binding buffer supplemented with 1 μg poly(dI-dC) at 4°C for 2 h with rotation. DNA-protein complexes were captured using streptavidin magnetic beads for 1 h at 4°C, followed by five washes and immunoblotting analyses.

### Immunoprecipitation

The transfected intact cells or enriched SCVs were washed once with PBS and lysed in ice-cold Buffer A (25 mM Tris-HCl, pH 7.5, 150 mM NaCl, 10% glycerol, 1% Triton X-100) supplemented with protease inhibitors. The lysates were briefly sonicated (20W power, 3-sec pulses with 7-sec intervals for 5 cycles) to ensure complete disruption and reduce viscosity. After centrifugation to remove debris, the supernatants were pre-cleared and subjected to anti-Flag immunoprecipitation following standard procedures. The immunoprecipitates were washed four times with ice-cold wash buffer B (25 mM Tris-HCl, pH 7.5, 150 mM NaCl, 0.5% Triton X-100), then eluted and for subsequent immunoblotting or MS analyses.

### Immunofluorescence microscopy and image analysis

For immunofluorescence staining, transfected or infected cells grown on coverslips were fixed in 4% paraformaldehyde for 10 min, permeabilized with 0.2% Triton X-100 for 15 min, and blocked with 2% BSA for 30 min. After incubation with primary antibodies, samples were labeled with species-matched Alexa Fluor-conjugated secondary antibodies. The nuclei were counterstained with DAPI for 2 min. Confocal images were acquired using a confocal microscope (FV3000RS, Olympus), with identical acquisition settings maintained across compared samples. Quantitative colocalization analysis was performed using ImageJ, and the colocalization index was calculated from at least 30 cells derived from three independent experiments. All images were processed uniformly with background subtraction, and threshold adjustments were applied equally to the control and experimental groups.

### Dual RNA-seq analysis

RAW264.7 cells infected with *S*. Typhimurium SL1344 for 10 h were lysed in TRIzol (Invitrogen) for total RNA isolation. Dual RNA-seq was performed by Novogene (Beijing, China), with reads aligned to both murine and *S*. Typhimurium SL1344 genomes. Gene expression was quantified using FPKM values, with significant differential expression defined as –log_10_(*p*-value) >1.3. Pathway enrichment of these differential expression genes (DEGs) was performed by GO and KEGG analyses using the DAVID online tool (https://david.ncifcrf.gov/).

### Metabolomics Analysis

RAW264.7 macrophages were mock-infected or infected with *S*. Typhimurium for 10 h and were washed twice with ice-cold PBS and rapidly quenched in liquid nitrogen for metabolite preservation. Untargeted metabolomic profiling was performed by PANOMIX Biomedical Tech Co., Ltd (Suzhou, China) using ultra-high-performance LC-MS. Raw data were processed using Xcalibur 4.0 software (Thermo Fisher Scientific) for peak alignment, retention time correction, and feature extraction. Multivariate statistical analysis was performed in MetaboAnalyst 5.0. Metabolite identification was achieved by matching against standard references in the HMDB and METLIN databases (mass accuracy< 5 ppm). Pathway analysis was conducted by mapping significantly altered metabolites (*p*< 0.05, fold change> 1.5) to KEGG pathways. All experiments included three independent biological replicates with quality control samples.

### MS analyses

To identify the ubiquitinated-containing peptides of GLUT1, purified Flag-GLUT1 protein was subjected to digestion with trypsin, and the resulting peptides were separated on an EASY-nLC 1200 system (Thermo Fisher Scientific). The nano liquid chromatography gradient was as follows: 0%–8% B in 3 min, 8%–28% B in 42 min, 28%–38% B in 5 min, and 38%–100% B in 10 min (solvent A: 0.1% formic acid in water; solvent B: 80% CH3CN in 0.1% formic acid). Peptides eluted from the capillary column were applied directly onto a Q Exactive Plus mass spectrometer by electrospray (Thermo Fisher Scientific) for MS and MS/MS analyses. Mass spectrometry data were searched against the amino acid sequence of GLUT1 and were performed with cleavage specificity, allowing four miscleavage events. Mass spectrometry data were searched with the variable modifications of methionine oxidation, ubiquitin addition to lysine, and acetylation of protein N termini.

For identification of the GLUT1-binding protein, immunoprecipitates were separated using SDS-PAGE, fixed, and visualized after silver staining as recommended by the manufacturer. An entire lane of bands was excised and subjected to in-gel trypsin digestion and MS/MS detections as described above. Identification of proteins was carried out using the Proteome Discoverer 2.2 program. Mass spectrometry data were searched against the human proteomes depending on the samples with carbamidomethylation of cysteine set as a fixed modification. The precursor mass tolerance was set to 10 ppm, and the fragment mass tolerance was set to 0.02 Da. A maximum false discovery rate of 1.0% was set for protein and peptide identifications.

### Statistical analysis

Statistical data are shown as mean ± standard deviation (SD) from at least three independent replicates. The student’s unpaired *t*-test method was used to compare two experimental groups, and the one-way analysis of variance (ANOVA) method was used to compare multiple groups. A difference is considered significant as follows: \**p*<0.05, \*\**p*< 0.01.

## Author Contributions

K.M. and D.X. conceived and designed the study. K.M., X.F., L.S., Y.W., and L.W. designed and performed the functional experiments. K.M., Y.W., and J.X. conducted the bioinformatics analyses. J.Y., Y.H., and J.X. prepared experimental materials. K.M., X.F., W.C., and X.T. performed the animal infection experiments. K.M., D.X., and X.F. analyzed the data and wrote the manuscript. Y.D. provided critical reagents and technical support. All authors discussed the results, critically reviewed the manuscript, and approved the final version.

## Acknowledgments

We thank members of the central laboratory of Taihe Hospital for helpful discussions and technical assistance. We thank the members of the Biomedical Research Institute of the Hubei University of Medicine for assisting in Confocal microscope operations. We thank Prof. Lu Feng and Prof. Lingyan Jiang (Nankai University) for kindly providing the *Salmonella* Typhi Ty2 strain.

This work was supported by the National Natural Science Foundation of China (32200156), the Innovative Research Program for Graduates of Hubei University of Medicine (YC202573), the Hubei Provincial Natural Science Foundation (2025AFD201, 2025AFB566), and the Shiyan Science and Technology Bureau Foundation (24Y028, 24Y031).

## Data Availability Statement

The raw sequence data reported in this paper have been deposited in the Genome Sequence Archive in BIG Data Center, Beijing Institute of Genomics (BIG) (https://bigd.big.ac.cn/gsa). The mass spectrometry proteomics data have been deposited in the ProteomeXchange Consortium via the iProX partner repository.

Other data supporting the findings of this study and source data are included in the supplementary data and available upon reasonable request.

